# On the duality of pain and pleasure processing: Why two dimensions of valence may be better than one

**DOI:** 10.1101/2025.01.22.634365

**Authors:** Zack Dulberg, Jonathan D. Cohen

## Abstract

Reinforcement learning treats reward maximization as a single objective, such that pain avoidance is implicit in pleasure seeking. However, humans appear to have distinct neural systems for processing pain and pleasure. This paper investigates the computational advantages of this separation through grid-world experiments. We demonstrate that modular architectures employing distinct *max* and *min* operators for value propagation outperform monolithic models in non-stationary environments. This separation allows agents to simultaneously grow and shrink learned values without interference, enabling both efficient reward collection and punishment avoidance. Additionally, these separate systems can be dynamically arbitrated using a mood-like mechanism for rapid adaptation. Our results suggest that separate pain and pleasure systems may have evolved to enable safe and efficient learning in changing environments.

*Nature has placed mankind under the governance of two sovereign masters, pain and pleasure*

Jeremy Bentham

## 1 Introduction

Human behavior, to first approximation, is based on avoiding pain and seeking pleasure. This is a re- statement of Freud’s “pleasure principle”. In his words: “any given process originates in an unpleasant state of tension and thereupon determines for itself such a path that its ultimate issue coincides with a relaxation of this tension, i.e. with avoidance of pain or with production of pleasure” (Freud, 1920). The fact that pain and pleasure have distinct natures seems true subjectively, but it is not obvious why this should be the case. In fact, Freud’s quote contains an implicit conflict – pain avoidance and pleasure seeking (two dimensions) are both mapped onto a single dimension; the ‘relaxation of tension’. Why then is one dimension alone, i.e., ‘seeking pleasure’, not a sufficient description of motivated behavior? After all, choosing a more pleasurable option automatically entails the ‘avoidance’ of a less pleasurable one.

This is in fact how standard reinforcement learning conceptualizes the issue. Scalar rewards are 1- dimensional, and reward maximization is the only goal; seeking reward and avoiding punishment are just two ways of describing the same objective. Any discrete categorization of valence on top of that is therefore arbitrary - a threshold drawn at some magnitude of reward. However, scalar reward maximization seems at odds with our intuition (and evidence) that approach (pleasure) and avoid (pain) systems are distinct. The goal of this paper is to bridge this gap, by suggesting some computational advantages of having two separate systems driving motivated behavior: one that is sensitive to good future outcomes, and one that is sensitive to bad ones. We thereby explain that pain and pleasure appear distinct because there are normative advantages, particularly in non-stationary environments, to maintaining separate learning systems for each.

## 2 Background

First, we will review evidence for separate valuation systems in the brain, as well as some existing models that attempt to account for this. There is longstanding (Dickinson & Dearing, 2014; Gray, 1982) and recent (Tian et al., 2024) behavioral and neuroscientific evidence for opposed motivational systems of valence; one that activates behavior to approach appetitive stimuli, and another that inhibits behavior to avoid aversive stimuli. This basic division has been elaborated in a variety of ways since. Classically, reward prediction errors have been observed in brain regions like the ventral tegmental area (Schultz et al., 1997), and the neurotransmitter dopamine has been shown to promote appetitive learning (Cools et al., 2009; Frank et al., 2004). Separate aversive prediction errors have been observed within the striatum (Eldar, Hauser, et al., 2016; Seymour et al., 2007), amygdala (Yacubian et al., 2006), insula (d’Acremont et al., 2009; Hoskin & Talmi, 2023), periaqueductal gray (Roy et al., 2014), orbitofrontal cortex (Hindi Attar et al., 2012) and dorsal raphe nucleues (Berg et al., 2014). There is evidence that the neuromodulator serotonin is responsible for these aversive error signals (Cools et al., 2008; Moran et al., 2018) and for avoidance learning in general (Dayan & Huys, 2008; Tanaka et al., 2009). This has led to the suggestion that the serotonin system may act in opposition to the dopamine system (Daw et al., 2002; Den Ouden et al., 2013). The evidence is also mixed; serotonin has been shown to encode rewards, (Bromberg-Martin et al., 2010; Li et al., 2016), dopamine is involved in aversive learning (de Jong et al., 2019; Matsumoto & Hikosaka, 2009; Watabe-Uchida & Uchida, 2018), and overlapping brain regions can respond to both gains and losses (Tom et al., 2007).

Despite the overlap, substantial evidence exists for distinct encoding and processing of different valances in the brain. We are interested in *why* that might be; what is the computational advantage? From a normative perspective, we are interested in the algorithms that neural systems dedicated to processing rewards and punishments might implement. We will now describe some candidate learning algorithms that attempt to distinguish in some way between different valences (the term *valence* is left intentionally imprecise; it is the details of the algorithms that will specify what differences we are really talking about).

## 3 Existing models

First, we will briefly introduce the temporal difference (TD) learning algorithm in a single system, and then describe variations that attempt to model valence-dependent differences in processing. We focus on model-free Q-learning, given the evidence that the brain makes use of this kind of algorithm (Daw et al., 2005; Montague et al., 1996; Schultz et al., 1997), and because the way TD-updates propagate value through state-space will be key to explaining why two systems for value learning may be better than one. Eqs. 1 - 3 display the standard Q-learning algorithm and decision process (with arguments dropped s.t. *Q* = *Q*(*st, at*) and *Q^′^*= *Q*(*st*+1*, at*+1) where possible).

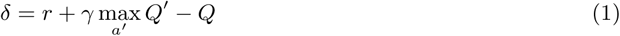

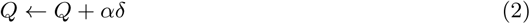

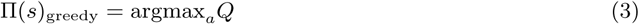

One variation of this algorithm makes the learning rate *α* a function of *δ* as in Eq. 4. This induces an asymmetry that is meant to represent risk-aversion (*k >* 0) or risk-tolerance (*k <* 0) (Mihatsch & Neuneier, 2002; Niv et al., 2012; Palminteri & Lebreton, 2022). Here, different valences are distinguished by the different rates of learning they induce.

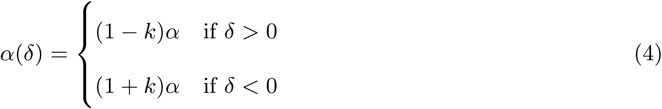

Another algorithmic variation of TD-learning is called *w*-pessimistic Q-learning. Rather than altering the learning rate, the error term itself is modified as in Eq. 5 such that best-case and worst-case scenarios are differentially weighted using an optimism parameter *w* (Dulberg et al., 2024; Gaskett, 2003; Zorowitz et al., 2020).

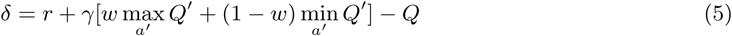

The parameter *w* controls the extent to which high vs. low values can propagate through the error term (i.e., max(*a, b*) is only a function of *a* when *a > b*). For example, having low *w* can be interpreted as an assumption that the world is not fully deterministic or controllable, and therefore the worst possible outcomes should play a role in learned valuations. Here, different valences are distinguished by which operator value predominantly propagates through into the temporal difference error (i.e., negatively-valenced pain stimuli would predominantly propagate through the min operator).

Both of these models still learn a single action value function, and therefore their action policy is uniquely determined by that function (i.e., Eq. 3 remains unchanged). An alternative approach is to learn two entirely separate value functions, similar to modular RL models (Amemori et al., 2011; Dulberg et al., 2023). For example, Enkhtaivan et al., 2020 defined ‘competing critics’ with two sets of Q-values, *Q*^+^ and *Q^−^*, updated as in Eqs. 6-7 with different biases toward risk, as in Eq. 8, such that a parameter *k^′^* determines how the learning rate differs for positive vs. negative prediction errors in each system. Then, *ɛ*-greedy action selection is performed after sampling a new set of Q-values from a uniform distribution between the two Q-value estimates for each action. This is a modularization of the risk-sensitive models just mentioned, and is therefore also a simple form of distributional RL, in which the agent learns two separate expectiles of future value are learned about (Dabney et al., 2020).

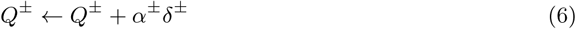

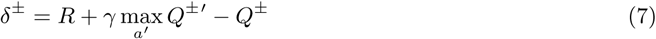

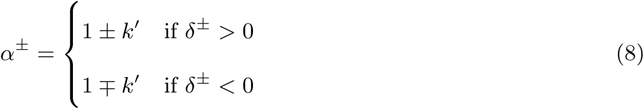

A final and influential model of multiple valuation systems is called MaxPain (Elfwing & Seymour, 2017), and is effectively a modularization of the *w*-pessimistic model defined in Eq. 5. This model splits the standard reward R into positive reward *r* and pain *p* components (Eq. 9), and then defines two Q-value functions, *Qr* that learns to maximize positive reward (Eq. 10), and *Qp* that learns to maximize pain (Eq. 11). For action selection, *Qr* and −*Qp* are linearly combined into *Qm*, with a mixing parameter *m* (Eq. 12). Finally, temporal difference errors are calculated using the best action according to *Qm* for reward (Eq. 13), but the worst action according to *Qm* for pain (Eq. 14).

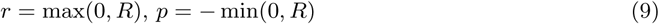

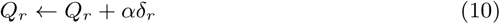

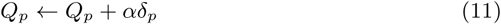

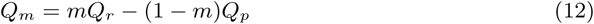

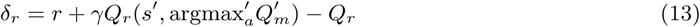

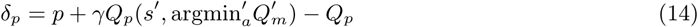

In effect, this algorithm attempts to maximize reward while minimizing pain (since the pain-maximizing Q-values are subtracted for the final policy). This algorithm has been extended to domains like deep learning and robotics (Wang & Uchibe, 2024; Wang et al., 2021), though with some details of the model varied, such as assuming the reward or pain maximizing systems control actions for value updates, as in Eqs. 15-16. Again, it should be noted for clarity, that using the max operator with pain *p* in Eq. 16 is equivalent to using the min operator with reward (i.e., −*p*).

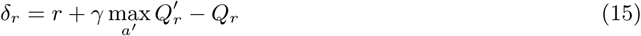

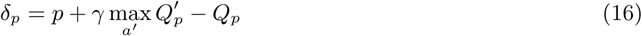

In summary, we have reviewed a variety of models that attempt to modify temporal difference learning to capture some kind of asymmetry between pain and pleasure. These included asymmetries between rewards, learning rates, and max/min operators within two broad classes of models: monolithic, in which a single value function is updated, and modular, in which two separate value functions are maintained. In the next section, we will test all these models to better understand their adaptive properties.

## 4 Experiment 1 - Remapping the value landscape

### 4.1 Approach

We build on a grid-world task described in Dulberg et al., 2024 designed to study grieving, in which a monolithic agent that learned using Eq. 5 had to adapt to a lost source of reward. In that study, when the optimism factor *w* was low (increasing the sensitivity of an agent to worst-case scenarios), negative rewards could be used to decrease the value of a lost object more quickly. Alternatively, it was also shown that high *w* more quickly *increased* value estimates associated with a lost source of punishment. Therefore, the *w* parameter promoted the propagation of positive value when high, and negative value when low. One can imagine situations in which each of these properties would be adaptive; with the gain of something negative or the loss of something positive, one would want to subtract from the value landscape, and with the gain of something positive or the loss of something negative, one would want to add to it. One can also imagine how the need to subtract value could be urgent, given the need to stay away from sources of danger, especially when the world is not fully controllable. But here we ask; what if both of these learning processes must take place at the same time? That is, how can an agent best avoid danger and maximize rewards in an environment where sources of both may be changing?

### 4.2 Environment

To answer this question, we constructed a grid-world task similar to the one used in Dulberg et al., 2024. The task is summarized in **Figure 1a**. The environment initially contained a positive reward in the top left of the grid in state *SA*, and a negative reward in the bottom right at state *SB*, and some walls that served to lengthen and delineate the pathway between them. The value map at *t* = *t^′^* shows what one selected Q-learning agent converged to after sufficient random exploration in this environment (this happens to be a MaxPain agent). Then, at time *t^′^*, reward and punishment locations swapped, with a more severe punishment replacing the previous reward (*rS_A_* = −5, representing a sudden increase in danger there). The agent was dropped at *SA*, and was given a finite horizon of *T* = 3000 steps to adapt its policy in order to avoid punishment and collect reward. It therefore had to overcome the behavioral policy induced by previous learning, which would move it away from *SB* and toward *SA*.

**Figure 1:**
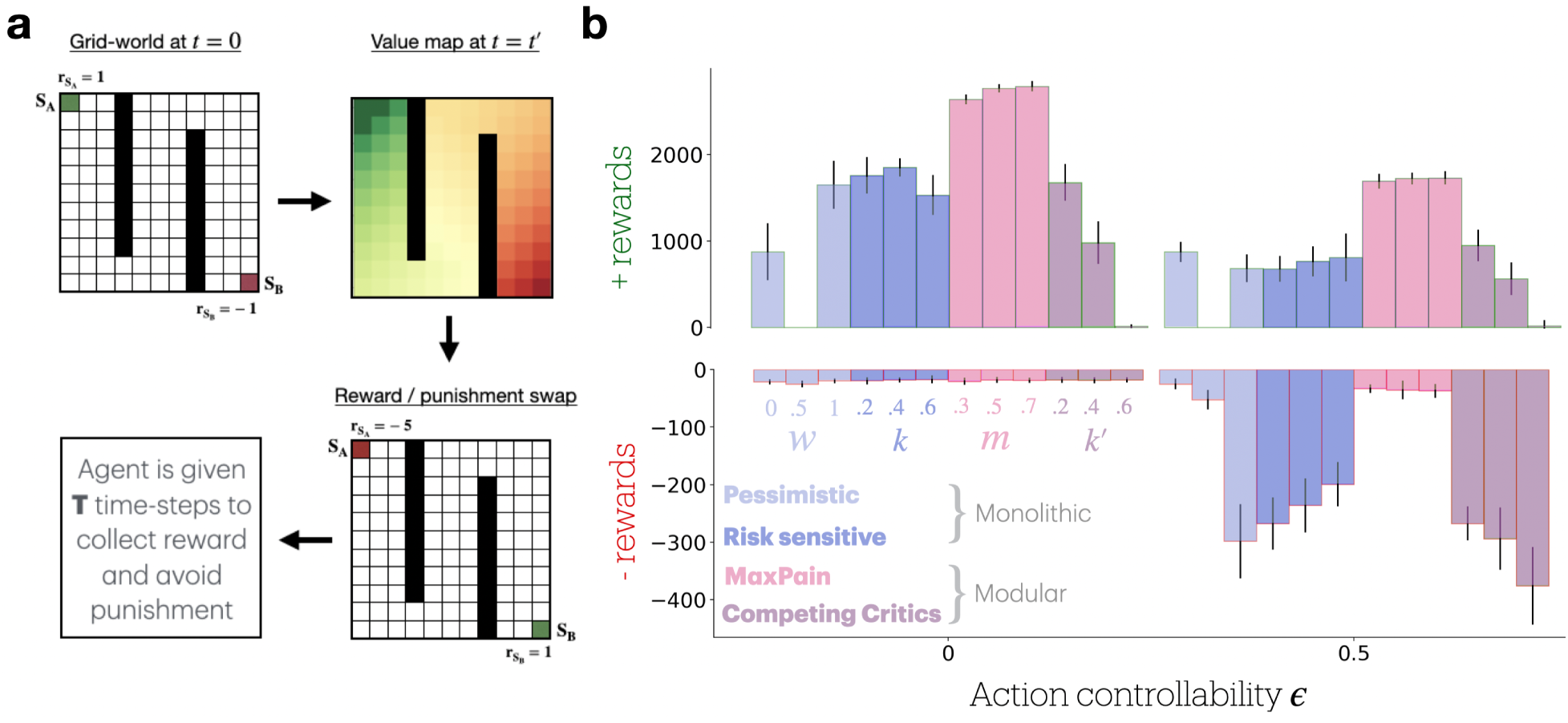
R**e**ward **/ punishment swap task. a:** Upper left: An 11x11 grid-world in which *S_A_* provided a reward *r_SA_* of 1 and *S_B_*provided a reward of *−*1. Upper right: A learned optimal state-value function (i.e. the maximum Q-value in each state after Q-values converged; high value states were brighter; MaxPain agent.). Lower right: Swap event when *r_SA_* was set to -5 and *r_SB_* was set to 1. Lower left: After the loss, the agent had *T* time-steps to learn a new policy, avoiding the new (and more severe) punishment in *S_A_*, and seeking positive reward from *S_B_*. **b:** Positive reward (top) and negative reward (bottom) collected by agents during *T* = 5000 time-steps after the swap event. These are shown for fully controllable actions (*ɛ* = 0) and when actions were only partially controllable (*ɛ* = 0.5), and each agent variant was tested for 3 parameter settings. Monolithic agents included *w*-pessimistic and risk-sensitive models, and modular agents included MaxPain and competing critics models (described in main text). The Maxpain model collected the most positive reward (was the most efficient), and suffered the least punishment when actions were partially controllable (was the safest). Bars show *µ ± σ* for *N* = 20 runs.

To speed up its learning, as in Dulberg et al., 2024, we allowed the agent to use DYNA memory replay (Sutton, 1991). In this case, we simply re-labelled the changed states with their new reward values (*rS_A_* = −5*, rS_B_* = 1) in the environment model used to generate replay transitions, i.e., providing the agent with knowledge about the changes in its environment. As in Dulberg et al., 2024, a replay ratio (*RR*) parameter determined how many transitions were replayed on each step in the environment, and *p_dwell_* determined what fraction of memories replayed were transitions into states that had changed. Default parameters used in these simulations are summarized in Table 1.

**Table 1:**
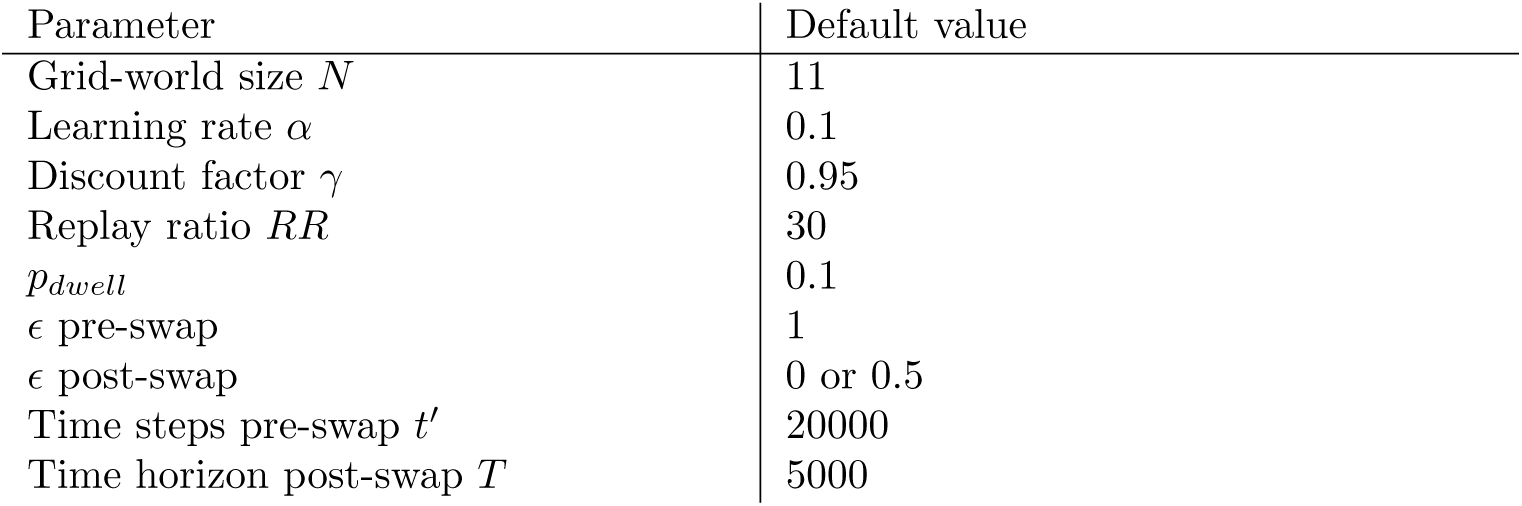
Default parameters used in current simulations if not otherwise specified.

### 4.3 Models

Since our goal was to understand why distinguishing between different valences might be adaptive, we tested the different ways of doing so enumerated above. We tested two model variants in the monolithic approach. The first, which we call *w***-pessimistic** Q-learning (Gaskett, 2003), is summarized in Eq. 5. We tested it with different values of its weight parameter *w*; lower *w* corresponding to greater pessimism with more weight on the min operator in TD-updates. The second model, which we call **risk-sensitive** (Mihatsch & Neuneier, 2002), is summarized in Eq. 4, and was tested with different values of the learning rate asymmetry parameter *k*; higher *k* corresponding to greater weight on negative prediction errors in TD-updates. We also tested two modular versions of these approaches. The first, referred to as **MaxPain** (Elfwing & Seymour, 2017), was tested for different mixing weights *m* as in Eq. 12; here, higher *m* indicated greater weight at the time of decision making on the Q-function that learned with the max operator. The second, referred to as **competing critics** (Enkhtaivan et al., 2020), was tested for different values of parameter *k^′^*; greater *k^′^* corresponded to a greater difference in risk sensitivity between separate Q-functions (in this model, one Q-function was risk-sensitive, and the other was risk-tolerant). In total, these 4 models were tested for 3 parameters settings each, and results reported as *µ* ± *σ* over *N* = 20 runs. We also studied two settings of the *ɛ* parameter (*ɛ* = [0, 0.5]) used for *ɛ*-greedy action selection after the swap at time *t^′^*. This was to simulate differences in the controllability of the agent; intuitively, it would be important to quickly escape the edge of a cliff if tripping off it could not be prevented 100% of the time. Therefore, we predicted that the benefits of any bias towards risk aversion or avoidance would be most prominent when *ɛ >* 0. When examining agent performance, we report total positive reward and total negative reward separately, to distinguish between efficiency (reward collection) and safety (punishment avoidance).

## 5 Results

### 5.1 Testing models on reward-punishment reversal task

Figure 1b summarizes the performance of all models across different parameter conditions. Monolithic agents that learned a single Q-value function included *w***-pessimistic** models for *w* ∈ (0, 0.5, 1) (which reflected pessimistic, neutral, and optimistic biases respectively) and **risk sensitive** models for *k* ∈ (0.2, 0.4, 0.6) (which reflected increasing levels of risk sensitivity). Modular agents that learned two Q-value functions included **MaxPain** models for *m* ∈ (0.3, 0.5, 0.7) (which reflected pessimistic, neutral, and optimistic Q-function mixtures respectively) and **competing critics** models for *k^′^* ∈ (0.2, 0.4, 0.6) (which reflected increasing levels of assymetry between risk-sensitive and risk-tolerant Q-functions). These models were tested under two action controllability conditions, *ɛ* = 0 (greedy) and *ɛ* = 0.5 (50% of actions chosen randomly). Accumulated rewards were split into total positive rewards (top) and total negative rewards (bottom).

We will summarize the performance of each set of models in turn. For the *w*-pessimistic models, in both controllability conditions, *w* = 0.5 was unable to collect any positive reward, in contrast to both *w* = 0 and *w* = 1. When actions were fully controllable, these models (along with all others) only accumulated a small amount of negative reward. However, when 50% of actions were random, greater *w* incurred greater punishment. This is because optimistically-biased agents stayed closer to the negative reward for longer, increasing the chance of randomly moving to it. The risk sensitive models were able to mitigate this effect only slightly; increasing *k* reduced negative reward to a small degree. These models also accumulated around the same amount of positive reward as the *w* = 1 model in both conditions. For the competing critics models, *k^′^* = 0.2 performed similarly to the best monolithic models, but increasing the asymmetry *k^′^* only worsened performance across all conditions. The best performing model overall was the modular MaxPain agent. Regardless of a bias *m* toward the optimistic or pessimistic policy in the final weighting, these models were able to collect more positive reward in both conditions, while simultaneously avoiding negative reward when actions were less controllable. In summary, the MaxPain agent was consistently the safest *and* the most efficient model in this task.

### 5.2 What factors account for the success of the MaxPain agent?

The MaxPain agent was best able to collect reward and avoid punishment after rewards and punishments swapped locations. What gave this modular model an advantage in the context of this non-stationarity? In this task, good performance required decreasing previously learned value estimates in the top left of the grid, while simultaneously increasing those in the bottom right of the grid. Naively, one might have assumed that the monolithic *w*-pessimistic model could also accomplish this when the max and min operator are both used as in *w* = 0.5. However, Figure 2a displays the value maps that result from the *w*-pessimistic model compared with the MaxPain model just before and 5k steps after the swap. The key observation here is that gradients of value did not propagate to the middle of the path for the *w*-pessimistic model. While the max and min operators allowed both high and low value terms to enter the error equation, they can also cancelled out when summed together. Only the MaxPain agent, that propagated high and low value terms independently in separate systems before combining them, could effectively alter the value landscape in both positive and negative directions at the same time.

**Figure 2:**
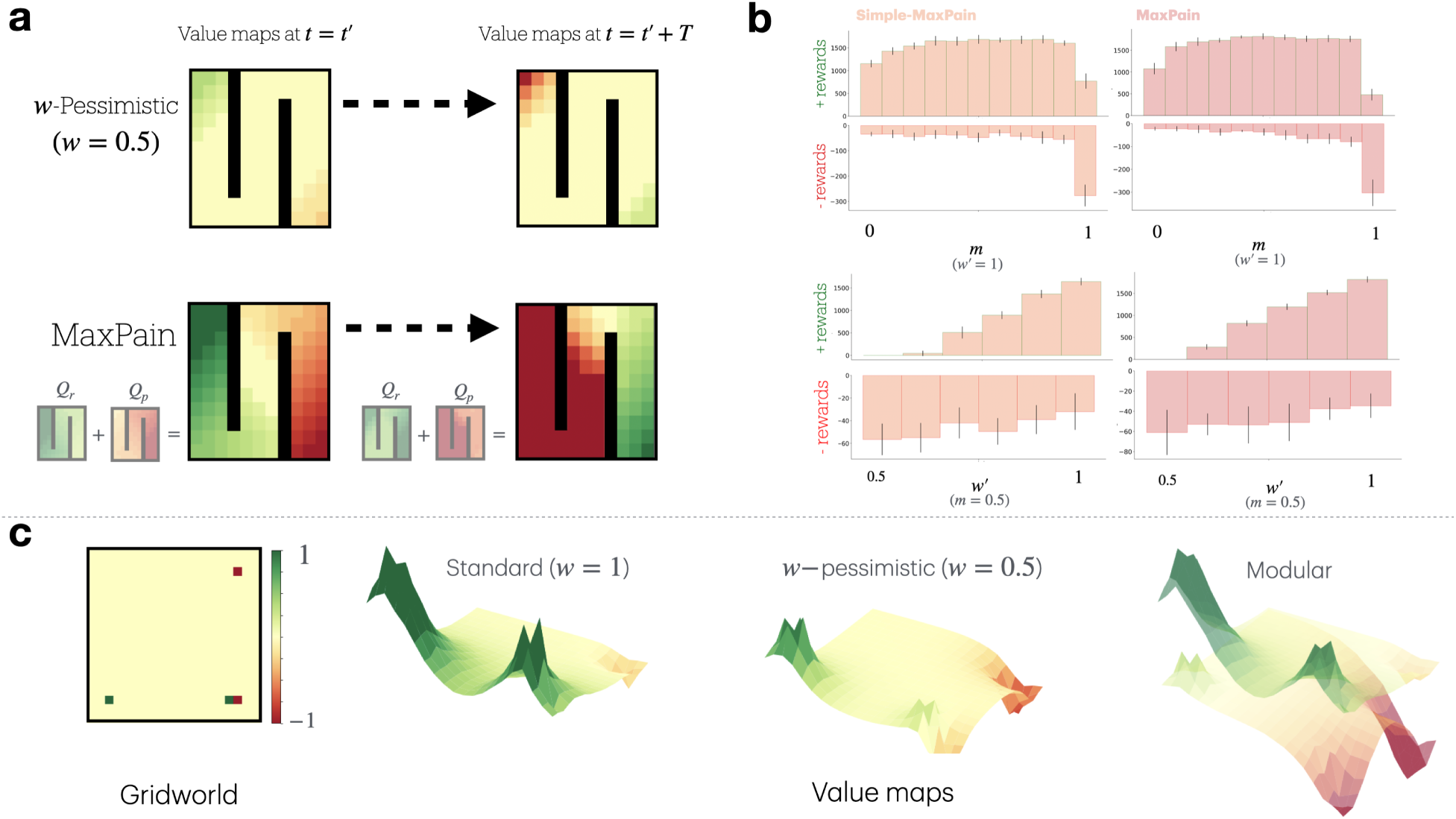
E**x**plaining **the advantage of the MaxPain model. a:** Value maps for the w-Pessimistic (w = 0.5) and MaxPain models before (left) and 5k steps after (right) the reward swap. The MaxPain model showed more effective alteration of the value landscape in both positive and negative directions simultane- ously, and gradients of value reachd deeper into the maze. **b:** Positive and negative rewards collected by simple-MaxPain (left) and MaxPain (right) models under different parameterizations. Top panels show the effect of varying the policy mixing parameter *m* for fully modular models (*w^′^* = 1). Bottom panels display the impact of varying *w^′^* in hybrid models for *m* = 0.5, which controlled the level of mixing max and min operators within modules. **c:** Visualization of learned value maps in an idealized grid-world (left) with one reward location, one punishment location, and one location with adjacent reward and punishment, for a stan- dard TD-agent (*w* = 1), a w-pessimistic agent (*w* = 0.5), and a modular simple-MaxPain agent. Interference was most obvious when comparing the poorly differentiated value-map at the adjacent reward/punishment location for the *w* = 0.5 agent with that of the modular agent.

We now ask, what were the crucial components of the MaxPain algorithm that allowed for this property, and how robust was the agent to different model parameterizations? Regarding the first question, the original model made a couple assumptions; the first was that the separate systems learned based on a disjoint set of rewards; the pleasure system learned only from positive rewards, while the pain system learned only from negative rewards (Eq. 9). We could simplify this assumption by allowing both systems to learn from all rewards, both positive and negative. Next, MaxPain estimated expected future value using the resultant *Qm*-function, such that the separate *Q*+ and *Q*− functions were learned off-policy. If one really wanted to learn to ‘maximize pain’ however (such that learning was truly sensitive to worst-possible scenarios), the *Q*− value function should learn as if it could control actions itself. This would also make computations more local, since learning could occur without access to the combined *Qm* policy. We therefore tested a model we refer to as **simple-MaxPain** that incorporated both these changes (resembling previously summarized Eqs. 15-16, but with *r* = *R* = −*p*).

We tested both variants of MaxPain agents side-by-side for different model parameterizations. First, we varied the mixing parameter *m*, assessing task performance for different relative weightings of the pain and pleasure module in the resultant value function *Qm*. Next, holding *m* constant at 0.5 (equal mixing), we relaxed the assumption that separate modules were exclusively optimistic or pessimistic, and made hybrid models that incorporated a *w*−pessimistic weighting inside each module. We varied a new parameter *w^′^* for both model variants, which represented how unequally optimism and pessimism were allocated between modules (where *w^′^* = 1 reduced to the original models in which max and min operators were exclusive to separate modules, and *w^′^* = 0.5 reduced to the monolithic case, i.e., both modules being identical). The relaxed simple-MaxPain model is summarized by the set of Eqs. 17-18. The relaxed MaxPain model incorporated the *w^′^* parameter in the same way (equations not shown).

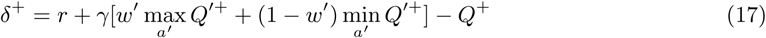

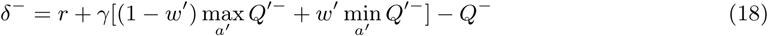

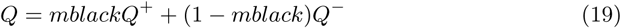

Figure 2b (top panels) first shows the task performance of fully modular (i.e., *w^′^* = 1) simple- MaxPain and MaxPain models for action controllability *ɛ* = 0.5. For both variants, performance was quite robust to *m*, only falling off at the extremes (when the agent used *Q^−^* or *Q*^+^ alone to make decisions). Basically, any other weighting of the two systems was sufficient to collect positive reward, and only when *Q*^+^ was used alone (*m* = 1) did the agents fail to avoid significant punishment. Next, Figure 2b (bottom panels) shows the effect of varying *w^′^* in the relaxed/hybrid variants. The result here was that any amount of within-module mixing hurt performance, decreasing positive reward and increasing punishment. Lastly, performance was basically identical between MaxPain and simple-MaxPain, suggesting that learning from disjoint reward sets, and combining component Q-functions before calculating future value, were not necessary to produce the properties that led to success in this task (the effect was the same when only one of these properties were changed at a time; this data was not presented to minimize redundancy).

To highlight the key properties of the relevant monolithic and modular agents, we provide an additional visualization of their learned value maps given an idealized grid-world in Figure 2c. In this gridworld, there were rewards of +1 alone (bottom-left), -1 alone (top-right), and +1 next to -1 (bottom-right). The learned value maps of each agent are shown to the right. The standard TD-agent (left, *w* = 1) learned green peaks of value wherever there was positive reward, and a modest red valley where there was isolated punishment. With balanced optimism and pessimism at *w* = 0.5 (middle), isolated peaks and valleys were less steep, and there was also interference: less distinction where reward and punishment were adjacent, and less propagation of reward and punishment into the middle of the value landscape. The modular agent (right, corresponding to simple-MaxPain) learned two separate value maps; one with prominent green peaks that propagated well from sources of reward, and one with prominent red valleys that propagated well from sources of punishment.

To summarize these results, the key feature that led to success in these modular agents appears to be the simultaneous use of max and min operators, given similar behavior between simple-MaxPain and MaxPain agents. Performance depended especially on the complete separation of these operators within-module, which helped avoid interference in value propagation. Then, the relative weighting of the two modules when combined to produce the resultant *Qm*-function was less important, as long as both modules had some minimum weight.

## 6 Experiment 2 - Mood as a policy arbitrator

Having established the potential usefulness of having separate value systems for both safety and efficiency during learning, we now consider an additional benefit of this configuration: Having two value estimates, and therefore two policies, can provide additional flexibility on timescales faster than gradual TD-learning can accommodate. That is, dynamically arbitrating between the two policies can allow an agent to adapt quickly to sudden changes in the environment. To be concrete, the visualization in Figure 2c for the modular agent consists of two separate value maps (one with green peaks, and one with red valleys) that could be used preferentially to seek reward or avoid punishment depending on the agent’s needs or the environmental context.

We explored one way to do this arbitration, according to the idea that the prediction error signal itself can provide additional information about the environmental context. Positive errors signal that conditions are better than expected, and negative errors signal that conditions are worse than expected. The accumulation of these errors over time has shown to correlate with mood empirically (Rutledge et al., 2014) and therefore has been used as a model of mood (Bennett et al., 2022; Dulberg et al., 2024; Eldar, Rutledge, et al., 2016; Emanuel & Eldar, 2023) that indicates how optimistic or pessimistic an agent should be about their environment. In Dulberg et al., 2024, this idea was used to simply measure expected mood trajectories in agents after a loss. Here, we used mood adaptively to arbitrate between pessimistic and optimistic policies in the modular agent. In that way, the agent was able to switch between optimistic and pessimistic behavioral policies quickly in response to environmental changes. We defined mood as in Eq. 20 as the accumulation of errors *δ* at a rate *η_mood_*, and used it in the weight factor when calculating the resultant Q-function *Qm*.

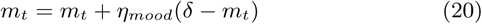

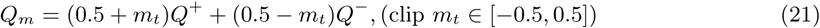

To demonstrate this mood-based adaptability, we constructed a task comprised of two stages, inspired by analogy to human psychological development. In the first stage, which we term development, the agent explored (until Q-value convergence) a grid-world in which reward was 0 everywhere except for adjacent rewards of 1 in the top left of the grid (state *SA*) and −1 in the state right next to *SA* (representing an area in state-space, like an attachment figure, that provided both positive and negative reinforcement). In the second stage, which we term adaptation / adult phase, we allowed the agent to act greedily in the grid-world, with *ɛ* = 0, learning turned off (*α* = 0), and mood initialized to 0.5 and then updated at a rate *η_mood_* = 0.01.

We first performed a proof-of-concept simulation, such that during the first 100 steps of the adaptation phase, *rS_A_* was set to 1, and during 100 steps after that *rS_A_* was set to −1. This reflected the same distribution of developmental reward magnitudes, but coming from the same state *SA*, instead of from adjacent states. This way, the agent would be surprised when *rS_A_* became negative. The agent started in state *SA*; perfect adaptation would therefore be to remain in *SA* for 100 steps, and then to avoid *SA* for 100 steps. We tested fixed and dynamic versions of earlier models on this task. Fixed models included the monolithic *w*-pessimistic model for *w* ∈ [0, 0.5, 1], and the modular MaxPain model for *m* ∈ [0.3, 0.5, 0.7].

The dynamic versions of these models replaced *w* and *m* parameters with a weight derived from the agent mood *mt*. Similar adaptive versions of the *w*-pessimistic (Karimpanal et al., 2023) and MaxPain (Mahajan et al., 2024) models have been previously explored, which also modulated time-varying *w* or *m* parameters using running accumulations of prediction errors. The difference between these two dynamic models is whether pessimism was modulated at the level of learning (TD-learning) (adaptive *w*) or at the level of decision making (policy mixing) (adaptive *m*).

Figure 3a shows the average performance of fixed and dynamic models on our adaptation task. Not surprisingly, fixed models could not adapt in time. According to their bias, they either only approached *SA* (*w, m >* 0.5), or only avoided it, without enough time to learn a new behavior in the face of a change. The dynamic model that modulated *w* showed similar results. Only the dynamic modular model (red) could adapt almost immediately, given its capacity to transition quickly between already-learned policies for approaching and avoiding the top-left area of the grid. It was therefore able to collect the most positive reward while avoiding the most punishment. This was a property unique to this model; despite this model and its modules learning from the same set of developmental data, it was able to differentiate separate approach and avoid policies given its ability to learn best and worst value functions in parallel. Finally, focusing just on the adaptive *m* model (dynamic modular agent), we explored the psychodynamic idea that a combination of agent make-up with its developmental environment (nature + nurture) might bias behavior to be more or less adaptive in different future environments. Despite learning that *w^′^* = 1 (no mixing of max and min operators within modules) worked best for the particular task studied in Figure 2, one still might expect this parameter to vary between individuals, given the evidence reviewed earlier suggesting pain and pleasure processing pathways do overlap. More generally, the benefits of modularity revealed here likely trade-off with other potential costs we did not study (such as increased conflict or decreased capacity to integrate information between modules).

**Figure 3:**
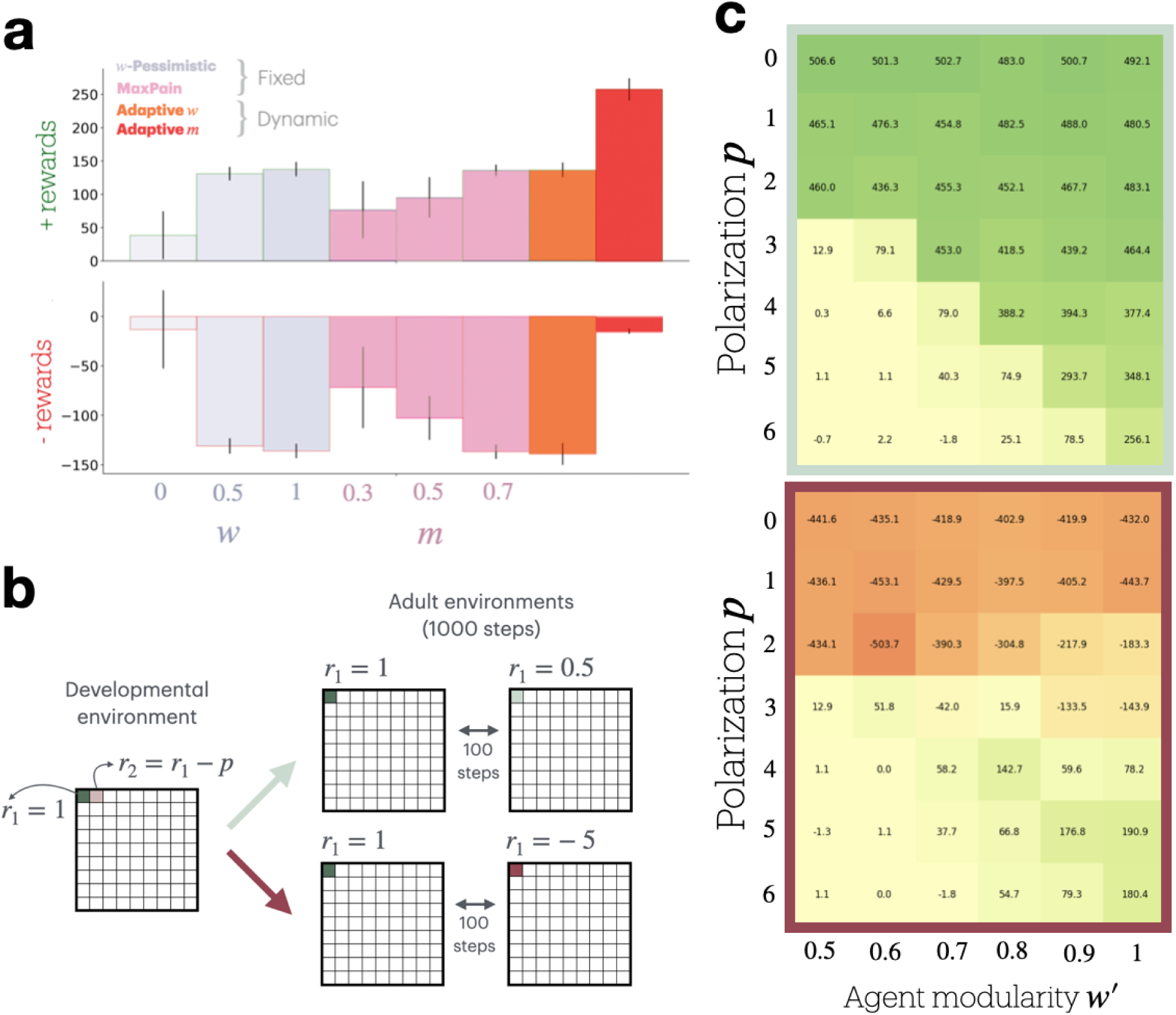
M**o**od **as a policy arbitrator. a:** Performance comparison of fixed and dynamic models in the adaptation task. The graph shows positive and negative rewards collected by different agent types over 200 steps (with reward changing from 1 to -1 at step 100). Fixed models (w-Pessimistic and MaxPain) and the dynamic w model showed limited adaptability, while the dynamic modular model (Adaptive *m*, red) demonstrated superior performance by quickly transitioning between approach and avoid policies. **b:** Experimental setup for testing the interaction between agent modularity and developmental environment. It illustrates the transition from a developmental environment with fixed rewards (*r*_1_ = 1*, r*_2_ = *r*_1_ *− p*) to two types of adult environments (1000 steps each). In the adult phase, *r*_1_ alternated every 100 steps between 1 and 0.5 (top) or between 1 and -5 (bottom), while *r*_2_ was set to 0. **c:** Heat maps showing total reward for different combinations of developmental environment polarization (*p*, y-axis) and agent modularity (*w^′^*, x-axis) in less polarized (top) and more polarized (bottom) adult environments. Colors reflect total reward magnitude.

We therefore constructed one more task variant, outlined in Figure 3b, in which the developmental phase contained one fixed reward in the top left location (*r*1 = 1), and a lesser reward *r*2 = *r*1 − *p* right next to it. Two dimensions were varied: The polarization of the agent’s modularity given by the parameter *w^′^* ∈ [0.5, 0.6, 0.7, 0.8, 0.9, 1] (nature), and the polarization of the developmental environment given by the parameter *p* ∈ [0, 1, 2, 3, 4, 5, 6] (nurture). Then, in the adaptation phase, learning was turned off (*α* = 0) and the agent acted greedily (*ɛ* = 0). In this phase, *r*2 = 0 (the adjacent punishment was removed, assumed to only have been present in the developmental environment), and *r*1 was again initialized to 1. However, *r*1 was made to alternate every 100 steps between 1 and 0.5 (top) or between 1 and −5 (bottom). To succeed in this environment, the agent needed to collect *r*1 for the full testing duration when it was consistently positive (top), but alternate approaching and avoiding it when it alternated with punishment (bottom). Because this phase contained no learning, we also added a term for mood homeostasis, as in Eq. 22, with *η_mood_*= 0.01, *η*_homeo_ = 0.02, and *m*_set-point_ = 0.5. This was done to give the agent an optimistic bias (Lefebvre et al., 2017), so it would return to seeking reward once enough time passed after avoidance. There is evidence for similar homeostatic mood dynamics in humans (Taquet et al., 2016).

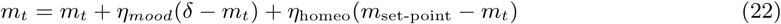

The analogy here again was that the agent, in its adult environment, encountered a source of reward similar to that from its developmental environment (i.e., a new relationship). However, this new reward source violated expectations half the time, either by providing only half the expected reward, or by providing punishment. The idea was to test how the interaction of agent and developmental polarization affected agent behavior in adult environments that were more or less polarized. Intuitively, being able to quickly transition between approach and avoid policies would be helpful in an environment where rewards alternated with punishment. However, strong avoidance behavior might be maladaptive when the new environment merely transitioned between a high and low level of reward. The clinical inspiration here was the phenomena of splitting, which is characterized by polarized object valuations (Story et al., 2024).

Results for this experiment are shown in Figure 3c. Total reward is displayed for the two dimensions of environment and agent polarization in the developmental phase, for less (top) and more (bottom) polarized adult / adaptation environments. Starting with the less polarized adult environment in which *r*1 alternated between 1 and 0.5, we see broadly that it was better to have a less polarized developmental environment. This was because the best strategy was to consistently collect *r*1. As developmental polarization increased, the agent’s pessimistic module was biased toward avoidance. Then, when negative reward prediction errors decreased mood, avoidance behavior was triggered. However, increasing modularity ameliorated this, by maintaining an approach module despite the developmental punishments. In less polarized developmental environments, the agent could also afford to be less modular for this reason. In contrast, in the polarized adult environment, the best performing agents were those from the polarized developmental environment, that were also modular enough to harness that polarity so they could both approach and avoid adaptively. When developmental polarization was low, the agent failed to avoid punishments, and when it was high but modularity was low, the agent failed to collect reward. In sum, agent modularity appeared to be generally adaptive, in that it allowed for harnessing, or buffering against, the effects of a polarized developmental environment. Developmental polarization was adaptive when it better reflected the structure of the adult testing environment. This experiment therefore suggests that the developmental environment may be a primary driver of adaptive or mal-adaptive behaviors (with the limiting assumption that developmentally learned value maps do not change), while the architecture of the agent can amplify or mitigate the impact of development. Splitting can therefore be interpreted as being adaptive for volatile environments, driven primarily by reward distributions in the developmental environment, but interacting with the intrinsic modularity of the agent.

## 7 Discussion

In this section, we asked the question: How can an adaptive agent safely and efficiently remap its learned value landscape? This question is specifically relevant to agents that learn model-free values, given the fact that agents with perfect world-models could just recalculate value maps if their environment changed. Since world-models are hard to learn and hard to compute, realistic agents also use model-free learning systems for their efficiency. However, this comes at the cost of inflexibility; gradually learned values must also be gradually unlearned to adapt to change. This motivates additional strategies to deal with this challenge. Here, we have suggested that having separate pain and pleasure systems is one such strategy.

Our simulations suggested that the best way to both grow and shrink a learned value landscape in parallel is to have separate systems that differ in the *operator*they use to propagate value. That is, the normative distinction between positive and negative valences is the way they are propagated through state-space during learning. We discovered this by studying the MaxPain architecture (Elfwing & Seymour, 2017), first showing it was safer and more efficient than competing models, and then noting these benefits relied on having isolated systems, one using only the *max* operator, and one using only the *min* operator, to estimate future value. A so-called safety-efficiency trade-off has been previously described, in which one is typically sacrificed for the other (Mahajan et al., 2024). However this is not always the case; in our task, safety and efficiency were aligned, since escaping dangerous regions more quickly also meant approaching rewarding regions more quickly. When actions were less controllable, escaping quickly was important to minimize punishment. In contrast, so-called risk-sensitive models incurred much more punishment in this scenario, simply because they could not propagate risk through their max operator.

Interestingly, the monolithic *w*-pessimistic model that balanced both the max and min operator in the prediction error term (i.e., *w*= 0.5), was the least efficient (collecting no reward). This was due to interference, such that positive and negative future values would cancel out in the error term. This model has been used to account for future actions being uncontrollable (Zorowitz et al., 2020), in which case the interference is a good thing; it inhibits approach if there is also danger in the same direction, reflecting an approach-avoid conflict (Epstein, 1978; Gray, 1982). However, this interference is not desirable when performance relies on simultaneously growing and shrinking value, or when approach and avoid are not in conflict. In this case, having separate systems allowed value efficiently propagate in both directions. This is reminiscent of separate representations being formed in the brain to avoid the interference that shared representations suffer from (Musslick et al., 2020). In this case, interference existed in the learning updates of a shared representation of value.

We also identified an additional benefit of modular value systems; the separate policies they induced could be used to respond to changes in the environment on a faster time-scale than learning, akin to a simple version of model-based revaluation (Dayan & Berridge, 2014). For example, we used the mood of the agent to dynamically adjust the contribution of each value module to the resultant Q-function used for decision making. In this way, when negative prediction errors accumulated, the agent shifted to using its pessimistic policy. Other models have also proposed that the RPE signal can dynamically adjust decision making based on environmental context (i.e., lean vs. rich resources) (Jaskir & Frank, 2023), and there is evidence that mood affects perception of outcomes (Eldar & Niv, 2015). Our account complements these, suggesting that mood may also arbitrate between policies that are distinguished by the operator used for TD-learning ^1^ (similar to how the accumulation of utility signals in the brain has been proposed to arbitrate between behavioral modes of exploration and exploitation (Aston-Jones & Cohen, 2005)). One could also view this from the perspective of latent cause inference, in which agents can learn that separate contexts should be treated differently, and can therefore learn value representations that depend on inferred latent causes (e.g., the presence of danger) (Gershman et al., 2015). One could consider the effective separation of pain and pleasure processing as a prior, encoded by evolution, that divides reality at a basic level into latent (and possibly overlapping) contexts for danger and opportunity, and mood as a simple model-based mechanism for context-dependent policy arbitration.

Although our goal was to understand valence at a computational level, some of our results may also bear on neuroanatomical questions. In attempting to simplify the MaxPain model, we noticed that one major modeling assumption, namely that positive rewards are sent to a pleasure system, and negative rewards are sent to a pain system, was not necessary for performance benefits. This is consistent with evidence that pain and reward signals overlap significantly in the brain (Leknes & Tracey, 2008). Still, it is an open question to what extent reward and punishment projections are distinct in the brain, and it is possible that how those signals interact with learning processes is a more distinguishing factor than their anatomical distributions (Combrisson et al., 2024).

Finally, there are a few clinically relevant considerations to note. First, one feature of clinical depression is reward insensitivity (Alloy et al., 2016). A modular model of valence naturally accommodates this. As we have shown, a module using a min operator is indeed relatively insensitive to positive reward (see Figure 2c; modular; bottom surface). Therefore, all it takes to exhibit reward insensitivity is the persistent activation of the pessimistic module (which, if our account of mood is correct, would be expected in the context of persistently low mood) (Shankman et al., 2007). These mood and policy dynamics, and any pathological attractors they produce, could potentially offer a formal account of the various emotional dynamics observed in depression (van Genugten et al., 2021).

Next, it has been suggested that anxiety is a disorder of planning (Sharp, 2024). That is, anxiety biases mental simulations (policy roll-outs) toward worst-case scenarios in order to avoid punishing future paths. One possible extension of our model is that modular policies could be used to simulate roll-outs in model-based RL. This may explain why some humans have such a strong tendency to seek out and ruminate on unpleasant mental experiences (Nolen-Hoeksema et al., 2008). If one considers a TD(*λ*) roll-out using the min-operator, this would lead to sequences that trace worst-case paths into the future. While worst-case pruning has been suggested as a feature of model-based planning algorithms (Huys et al., 2012), and rumination as a model-based avoidance strategy (problem solving) (Bedder et al., 2024), we suggest that such sequences may not only lead to pruning or problem-solving, but also model-free value propagation through the min-operator of an aversion module. This would be consistent with other accounts of non-local replay to propagate reward-related information (Krausz et al., 2023). It may turn out that the mental approach of worst-case scenarios most effectively promotes behavioral avoidance, bringing to mind the phenomena of intrusive thoughts (Clark & Rhyno, 2005), and the quote by Alfred North Whitehead: “*The purpose of thinking is to let the ideas die instead of us dying.*”

Lastly, our results relate to psychological development, with splitting as an example. Splitting refers to subjects assigning qualities of good and bad in an extreme (”black and white”) manner, exhibiting both significant overvaluation and undervaluation, and sometimes rapidly alternating between the two (Gerson, 1984). Splitting has previously been modeled from a Bayesian perspective involving the inference of good and bad latent causes (Story et al., 2024). Our work suggests that splitting dynamics could emerge from interactions between the statistics of the developmental environment, the modularity of the agent architecture (possibly itself learned from statistics (Giallanza et al., 2024)), and the emergent mood dynamics that might arbitrate between optimistic and pessimistic behavioral policies. Such behaviors could turn out to be adaptive or maladaptive depending on how closely future environments resemble developmental ones. However, our experiment should be viewed as a limited proof-of-concept for studying phenomena like splitting and agent-development interactions; the dynamics of splitting are likely multi- factorial, and should be studied further, over a wider set of parameters, in a variety of more complex environments that also consider interactions between learning and mood.

## 8 Conclusion

This study explored the computational advantages of separate pain and pleasure systems in reinforcement learning agents. We demonstrated that modular architectures, particularly those employing distinct max and min operators for value propagation, excel at efficiently remapping value landscapes in non-stationary environments. This approach allows agents to simultaneously grow and shrink learned values, overcoming interference issues present in monolithic models. Furthermore, we showed how these separate systems can be dynamically arbitrated using a mood-like mechanism, enabling rapid adaptation to environmental changes. Our simulations also highlighted the interplay between agent architecture and developmental environment in shaping adaptive behaviors. These findings offer insights into the potential evolutionary origins of distinct valence systems and provide a computational framework for understanding core psychodynamic phenomena related to depression, anxiety and personality.

## Footnotes

1 There may also be additional operators used other than max and min, as studied in Wang and Uchibe, 2024

## References

1. Alloy, L. B., Olino, T., Freed, R. D., & Nusslock, R. (2016). Role of reward sensitivity and processing in major depressive and bipolar spectrum disorders. Behavior therapy, 47 (5), 600–621.

2. Amemori, K.-i., Gibb, L. G., & Graybiel, A. M. (2011). Shifting responsibly: The importance of striatal modularity to reinforcement learning in uncertain environments. Frontiers in human neuroscience, 5, 47.

3. Aston-Jones, G., & Cohen, J. D. (2005). An integrative theory of locus coeruleus-norepinephrine function: Adaptive gain and optimal performance. Annu. Rev. Neurosci., 28, 403–450.

4. Bedder, R. L., Hitchcock, P. F., & Sharp, P. B. (2024). Unraveling repetitive negative thinking with reinforcement learning.

5. Bennett, D., Davidson, G., & Niv, Y. (2022). A model of mood as integrated advantage. Psychological Review, 129 (3), 513.

6. Berg, B. A., Schoenbaum, G., & McDannald, M. A. (2014). The dorsal raphe nucleus is integral to negative prediction errors in pavlovian fear. European Journal of Neuroscience, 40 (7), 3096–3101.

7. Bromberg-Martin, E. S., Hikosaka, O., & Nakamura, K. (2010). Coding of task reward value in the dorsal raphe nucleus. Journal of Neuroscience, 30 (18), 6262–6272.

8. Clark, D. A., & Rhyno, S. (2005). Unwanted intrusive thoughts in nonclinical individuals: Implications for clinical disorders.

9. Combrisson, E., Basanisi, R., Gueguen, M. C., Rheims, S., Kahane, P., Bastin, J., & Brovelli, A. (2024). Neural interactions in the human frontal cortex dissociate reward and punishment learning. Elife, 12, RP92938.

10. Cools, R., Frank, M. J., Gibbs, S. E., Miyakawa, A., Jagust, W., & D’Esposito, M. (2009). Striatal dopamine predicts outcome-specific reversal learning and its sensitivity to dopaminergic drug administration. Journal of Neuroscience, 29 (5), 1538–1543.

11. Cools, R., Robinson, O. J., & Sahakian, B. (2008). Acute tryptophan depletion in healthy volunteers enhances punishment prediction but does not affect reward prediction. Neuropsychopharmacology, 33 (9), 2291–2299.

12. Dabney, W., Kurth-Nelson, Z., Uchida, N., Starkweather, C. K., Hassabis, D., Munos, R., & Botvinick, M. (2020). A distributional code for value in dopamine-based reinforcement learning. Nature, 577 (7792), 671–675.

13. d’Acremont, M., Lu, Z.-L., Li, X., Van der Linden, M., & Bechara, A. (2009). Neural correlates of risk prediction error during reinforcement learning in humans. Neuroimage, 47 (4), 1929–1939.

14. Daw, N. D., Kakade, S., & Dayan, P. (2002). Opponent interactions between serotonin and dopamine. Neural networks, 15 (4-6), 603–616.

15. Daw, N. D., Niv, Y., & Dayan, P. (2005). Uncertainty-based competition between prefrontal and dorsolateral striatal systems for behavioral control. Nature neuroscience, 8 (12), 1704– 1711.

16. Dayan, P., & Berridge, K. C. (2014). Model-based and model-free pavlovian reward learning: Revaluation, revision, and revelation. *Cognitive, Affective*, & Behavioral Neuroscience, 14, 473–492.

17. Dayan, P., & Huys, Q. J. M. (2008). Serotonin, inhibition, and negative mood. PLoS computational biology, 4 (2), e4.

18. de Jong, J. W., Afjei, S. A., Dorocic, I. P., Peck, J. R., Liu, C., Kim, C. K., Tian, L., Deisseroth, K., & Lammel, S. (2019). A neural circuit mechanism for encoding aversive stimuli in the mesolimbic dopamine system. Neuron, 101 (1), 133–151.

19. Den Ouden, H. E., Daw, N. D., Fernandez, G., Elshout, J. A., Rijpkema, M., Hoogman, M., Franke, B., & Cools, R. (2013). Dissociable effects of dopamine and serotonin on reversal learning. Neuron, 80 (4), 1090–1100.

20. Dickinson, A., & Dearing, M. F. (2014). Appetitive—aversive interactions and inhibitory processes. In Mechanisms of learning and motivation (pp. 203–231). Psychology Press.

21. Dulberg, Z., Dubey, R., Berwian, I. M., & Cohen, J. D. (2023). Having multiple selves helps learning agents explore and adapt in complex changing worlds. Proceedings of the National Academy of Sciences, 120 (28), e2221180120.

22. Dulberg, Z., Dubey, R., & Cohen, J. D. (2024). Adapting to loss: A normative account of grief. bioRxiv, 2024–02.

23. Eldar, E., Hauser, T. U., Dayan, P., & Dolan, R. J. (2016). Striatal structure and function predict individual biases in learning to avoid pain. Proceedings of the National Academy of Sciences, 113 (17), 4812–4817.

24. Eldar, E., & Niv, Y. (2015). Interaction between emotional state and learning underlies mood instability. Nature communications, 6 (1), 6149.

25. Eldar, E., Rutledge, R. B., Dolan, R. J., & Niv, Y. (2016). Mood as representation of momentum. Trends in cognitive sciences, 20 (1), 15–24.

26. Elfwing, S., & Seymour, B. (2017). Parallel reward and punishment control in humans and robots: Safe reinforcement learning using the maxpain algorithm. 2017 *Joint IEEE International Conference on Development and Learning and Epigenetic Robotics (ICDL-EpiRob)*, 140– 147.

27. Emanuel, A., & Eldar, E. (2023). Emotions as computations. Neuroscience & Biobehavioral Reviews, 144, 104977.

28. Enkhtaivan, E., Nishimura, J., Ly, C., & Cochran, A. (2020). A competition of critics in human decision-making. *bioRxiv*.

29. Epstein, S. (1978). Avoidance–approach: The fifth basic conflict. Journal of Consulting and Clinical Psychology, 46 (5), 1016.

30. Frank, M. J., Seeberger, L. C., & O’reilly, R. C. (2004). By carrot or by stick: Cognitive reinforcement learning in parkinsonism. Science, 306 (5703), 1940–1943.

31. Freud, S. (1920). Beyond the pleasure principle. *London**: The*.

32. Gaskett, C. (2003). Reinforcement learning under circumstances beyond its control.

33. Gershman, S. J., Norman, K. A., & Niv, Y. (2015). Discovering latent causes in reinforcement learning. Current Opinion in Behavioral Sciences, 5, 43–50.

34. Gerson, M.-J. (1984). Splitting: The development of a measure. Journal of Clinical Psychology, 40 (1), 157–162.

35. Giallanza, T., Campbell, D., & Cohen, J. D. (2024). Toward the emergence of intelligent control: Episodic generalization and optimization. Open Mind, 8, 688–722.

36. Gray, J. A. (1982). Pŕecis of the neuropsychology of anxiety: An enquiry into the functions of the septo-hippocampal system. Behavioral and brain sciences, 5 (3), 469–484.

37. Hindi Attar, C., Finckh, B., & Büchel, C. (2012). The influence of serotonin on fear learning. Hoskin, R., & Talmi, D. (2023). Adaptive coding of pain prediction error in the anterior insula. European Journal of Pain, 27 (6), 766–778.

38. Huys, Q. J., Eshel, N., O’Nions, E., Sheridan, L., Dayan, P., & Roiser, J. P. (2012). Bonsai trees in your head: How the pavlovian system sculpts goal-directed choices by pruning decision trees. PLoS computational biology, 8 (3), e1002410.

39. Jaskir, A., & Frank, M. J. (2023). On the normative advantages of dopamine and striatal opponency for learning and choice. Elife, 12, e85107.

40. Karimpanal, T. G., Le, H., Abdolshah, M., Rana, S., Gupta, S., Tran, T., & Venkatesh, S. (2023). Balanced q-learning: Combining the influence of optimistic and pessimistic targets. Artificial Intelligence, 325, 104021.

41. Krausz, T. A., Comrie, A. E., Kahn, A. E., Frank, L. M., Daw, N. D., & Berke, J. D. (2023). Dual credit assignment processes underlie dopamine signals in a complex spatial environment. Neuron, 111 (21), 3465–3478.

42. Lefebvre, G., Lebreton, M., Meyniel, F., Bourgeois-Gironde, S., & Palminteri, S. (2017). Behavioural and neural characterization of optimistic reinforcement learning. Nature Human Behaviour, 1 (4), 0067.

43. Leknes, S., & Tracey, I. (2008). A common neurobiology for pain and pleasure. Nature reviews neuroscience, 9 (4), 314–320.

44. Li, Y., Zhong, W., Wang, D., Feng, Q., Liu, Z., Zhou, J., Jia, C., Hu, F., Zeng, J., Guo, Q., et al. (2016). Serotonin neurons in the dorsal raphe nucleus encode reward signals. Nature communications, 7 (1), 10503.

45. Mahajan, P., Tong, S., Lee, S. W., & Seymour, B. (2024). Balancing safety and efficiency in human decision making. bioRxiv, 2024–01.

46. Matsumoto, M., & Hikosaka, O. (2009). Two types of dopamine neuron distinctly convey positive and negative motivational signals. Nature, 459 (7248), 837–841.

47. Mihatsch, O., & Neuneier, R. (2002). Risk-sensitive reinforcement learning. Machine learning, 49, 267–290.

48. Montague, P. R., Dayan, P., & Sejnowski, T. J. (1996). A framework for mesencephalic dopamine systems based on predictive hebbian learning. Journal of neuroscience, 16 (5), 1936–1947.

49. Moran, R. J., Kishida, K. T., Lohrenz, T., Saez, I., Laxton, A. W., Witcher, M. R., Tatter, S. B., Ellis, T. L., Phillips, P. E., Dayan, P., et al. (2018). The protective action encoding of serotonin transients in the human brain. Neuropsychopharmacology, 43 (6), 1425–1435.

50. Musslick, S., Saxe, A., Hoskin, A. N., Sagiv, Y., Reichman, D., Petri, G., & Cohen, J. D. (2020). On the rational boundedness of cognitive control: Shared versus separated representa- tions.

51. Niv, Y., Edlund, J. A., Dayan, P., & O’Doherty, J. P. (2012). Neural prediction errors reveal a risk-sensitive reinforcement-learning process in the human brain. Journal of Neuroscience, 32 (2), 551–562.

52. Nolen-Hoeksema, S., Wisco, B. E., & Lyubomirsky, S. (2008). Rethinking rumination. Perspectives on psychological science, 3 (5), 400–424.

53. Palminteri, S., & Lebreton, M. (2022). The computational roots of positivity and confirmation biases in reinforcement learning. Trends in Cognitive Sciences, 26 (7), 607–621.

54. Roy, M., Shohamy, D., Daw, N., Jepma, M., Wimmer, G. E., & Wager, T. D. (2014). Representation of aversive prediction errors in the human periaqueductal gray. Nature neuroscience, 17 (11), 1607–1612.

55. Rutledge, R. B., Skandali, N., Dayan, P., & Dolan, R. J. (2014). A computational and neural model of momentary subjective well-being. Proceedings of the National Academy of Sciences, 111 (33), 12252–12257.

56. Schultz, W., Dayan, P., & Montague, P. R. (1997). A neural substrate of prediction and reward. Science, 275 (5306), 1593–1599.

57. Seymour, B., Daw, N., Dayan, P., Singer, T., & Dolan, R. (2007). Differential encoding of losses and gains in the human striatum. Journal of Neuroscience, 27 (18), 4826–4831.

58. Shankman, S. A., Klein, D. N., Tenke, C. E., & Bruder, G. E. (2007). Reward sensitivity in depression: A biobehavioral study. Journal of abnormal psychology, 116 (1), 95.

59. Sharp, P. B. (2024). Anxiety involves planning.

60. Story, G. W., Smith, R., Moutoussis, M., Berwian, I. M., Nolte, T., Bilek, E., Siegel, J. Z., & Dolan, R. J. (2024). A social inference model of idealization and devaluation. Psychological Review, 131 (3), 749.

61. Sutton, R. S. (1991). Dyna, an integrated architecture for learning, planning, and reacting. ACM Sigart Bulletin, 2 (4), 160–163.

62. Tanaka, S. C., Shishida, K., Schweighofer, N., Okamoto, Y., Yamawaki, S., & Doya, K. (2009). Serotonin affects association of aversive outcomes to past actions. Journal of Neuroscience, 29 (50), 15669–15674.

63. Taquet, M., Quoidbach, J., De Montjoye, Y.-A., Desseilles, M., & Gross, J. J. (2016). Hedonism and the choice of everyday activities. Proceedings of the national Academy of Sciences, 113 (35), 9769–9773.

64. Tian, Z., Song, J., Zhao, X., Zhou, Y., Chen, X., Le, Q., Wang, F., Ma, L., & Liu, X. (2024). The interhemispheric amygdala-accumbens circuit encodes negative valence in mice. Science, 386 (6722), eadp7520.

65. Tom, S. M., Fox, C. R., Trepel, C., & Poldrack, R. A. (2007). The neural basis of loss aversion in decision-making under risk. Science, 315 (5811), 515–518.

66. van Genugten, C. R., Schuurmans, J., van Ballegooijen, W., Hoogendoorn, A. W., Smit, J. H., & Riper, H. (2021). Discovering different profiles in the dynamics of depression based on real–time monitoring of mood: A first exploration. Internet Interventions, 26, 100437.

67. Wang, J., Elfwing, S., & Uchibe, E. (2021). Modular deep reinforcement learning from reward and punishment for robot navigation. Neural Networks, 135, 115–126.

68. Wang, J., & Uchibe, E. (2024). Reward-punishment reinforcement learning with maximum entropy. arXiv preprint arXiv:2405.11784.

69. Watabe-Uchida, M., & Uchida, N. (2018). Multiple dopamine systems: Weal and woe of dopamine. Cold spring harbor symposia on quantitative biology, 83, 83–95.

70. Yacubian, J., Glascher, J., Schroeder, K., Sommer, T., Braus, D. F., & Büchel, C. (2006). Dissociable systems for gain-and loss-related value predictions and errors of prediction in the human brain. Journal of Neuroscience, 26 (37), 9530–9537.

71. Zorowitz, S., Momennejad, I., & Daw, N. D. (2020). Anxiety, avoidance, and sequential evaluation. *Computational Psychiatry (Cambridge*, Mass.), 4.

